# Disruption in Patterning in the Whisker-to-Barrel Cortex Pathway Alters Behavior

**DOI:** 10.1101/2025.11.06.687024

**Authors:** Joanna M.F. Lutchman, Sarah A.F. Lutchman, Joshua C. Brumberg

**Author notes:** <u>Corresponding Author:</u> Joshua C. Brumberg, Department of Psychology, Queens College, CUNY, 65-30 Kissena Boulevard, Flushing, NY 11367. Joanna Lutchman –, Sarah Lutchman –.

## Abstract

Mice use their whiskers to convey sensory information, navigate, and explore their environment. We investigated the behavioral impact of the disruption of somatotopic patterning along the whisker-to-barrel pathway utilizing two mouse models: Barrelless (BRL) mice, an adenylyl cyclase 1 variation, in which somatotopic patterning is absent in the barrel cortex, and Prrxl1^−/−^mice, a genetic knockout in which patterning is disrupted along the entire lemniscal pathway. A textured novel object recognition test was conducted to investigate whisker-dependent discriminatory behavior, and an open field test was conducted to investigate exploratory behavior. Results were compared to an outbred strain (CD-1) and demonstrated that BRL mice were able to discriminate, whereas Prrxl1^−/−^ mice were unable to discriminate between textures, and that both strains exhibited increased anxiety. Exploratory and locomotor behavior increased in BRL mice but decreased in Prrxl1^−/−^ mice. Together, the results suggest that somatotopy may be related to behavioral phenotype.

## 1. Introduction

Mice rely heavily on their whiskers to navigate, explore, and survive in their environment. Whisking – the active movement of whiskers – transmits sensory information along the lemniscal and paralemniscal pathways to barrel structures in the primary somatosensory cortex (S1) [1, 2]. The patterning of the whiskers on the mystacial pad is maintained as a one-to-one representation along the whisker-to-barrel pathway as barrelettes in the trigeminal nuclei complex located in the brainstem, barreloids in the thalamus, and barrels in the S1 region of the somatosensory cortex [3]. Discovered by Woolsey and van der Loos in 1970, the cortical barrel field is located in layer IV of the cortex and comprises “barrels,” each corresponding to one whisker on the contralateral face [4].

In Barrelless (BRL) mice, a spontaneous mutation occurs in which the thalamocortical afferents travel to layer IV of the somatosensory cortex but do not segregate into barrels [5, 6]. This mutation is caused by the disruption of the adenylyl cyclase type I (Adcy1 or AC1) gene, which catalyzes the conversion of ATP to cyclic AMP. Cyclic AMP signaling is important for the patterning of somatosensory maps, resulting in defects in neuronal pattern formation [7]. The somatotopic patterning of the whisker-to-barrel pathway is disrupted only in the barrel cortex of BRL mice, while the patterning of barrelettes in the principal sensory nucleus (PrV) and barreloids in the thalamus are maintained [5–8].

Prrxl1 (previously known as DRG11) is a paired homeodomain transcription factor necessary for patterning during development [9, 10]. Prrxl1 is expressed in the PrV of the trigeminal nuclei [9, 11]. The deletion of the Prrxl1 gene results in disruption of somatotopic patterning in the whisker-to-barrel pathway with an absence of barrelettes in the PrV, barreloids in the thalamus, and barrels in the cortex, confirmed by cytochrome oxidase staining [10, 12].

In a previous study, Arakawa et al., [7] investigated the performance of BRL mice on various behavioral tasks, such as sensorimotor ability tests, discrimination tasks, and general motor ability tests. Control mice crossed a larger distance compared to BRL mice in the gap crossing test, which evaluated the mice’s ability to detect and judge distances using their whiskers. In the whisking test, which analyzed the frequency of whisker movements, BRL mice displayed a lower ratio of active whisking when in contact with an object compared to the controls. An object shape discrimination task and an object texture discrimination task were conducted to test the mouse’s ability to detect and remember different shapes as well as to assess whisker-mediated texture discrimination. The performance of the BRL mice did not differ on the object shape discrimination task compared to the controls; however, the BRL mice were not able to discriminate between familiar and unfamiliar textures in the texture discrimination task. An open field test was also conducted, but it only investigated the locomotor activity by measuring the number of lines crossed, not the general activity level. Additionally, only male mice were used; therefore, potential sex differences were not investigated.

Pain behavior and motor activity in Prrxl1^−/−^ mice were investigated and a reduction in the response of Prrxl1^−/−^ mice to thermal and chemical nociceptive stimuli, as well as mechanical stimulation were observed [9, 13]. However, sensorimotor function was assessed using a rotarod treadmill test, and no deficits in motor activity were reported [9]. Bakalar et al., [11] investigated ingestive behavior and reported impairment in feeding, including the handling and consumption of solid food. Although behavioral studies were conducted on both strains, there is a lack of reported information focusing on the effect of somatotopic pattern disruption on the exploratory and whisker-dependent discriminatory behaviors in these two strains of mice, and due to differences in experimental conditions, direct comparisons are not possible. We aim to examine the impact of somatotopic pattern disruption on exploratory and whisker-dependent discriminatory behavioral tasks, and to investigate any sex difference using male and female BRL and Prrxl1^−/−^ mice, neither of which were previously reported.

BRL and Prrxl1^−/−^ mice exhibit distinct disruptions in somatotopic representation along the whisker-to-barrel pathway [5–8, 10, 12]. Somatotopic patterning is abolished only in the barrel cortex of BRL mice, and the PrV of the lemniscal brainstem nucleus, thalamus, and barrel cortex in Prrxl1^−/−^ mice. As such, these two strains of mice can be used to study the behavioral effects of differential pattern disruption along the whisker-to-barrel pathway. This study aims to examine whisker-dependent texture discrimination behavior among the two different strains of mice using the textured novel object recognition test to assess the ability to discriminate between different textures, since more time is spent investigating a novel texture as opposed to a familiar texture in the absence of any behavioral impairment [14–16]. This study also aims to evaluate exploratory behavior using the open field behavioral test, since more time spent in the inner arena is associated with an increase in exploratory behavior [17–19]. Since sensory information from the environment is relayed along the whisker-to-barrel pathway, it is hypothesized that the disruption of patterning in the PrV and S1 region of this pathway will result in the behavioral impairment of BRL and Prrxl1^−/−^ mice on textured novel object recognition and open field behavioral tests.

## 2. Materials & Methods

### 2.1. Subjects

All experiments were approved by the Queens College Institutional Animal Care and Use Committee (IACUC protocol #208). A total of 153 mice were used in this study. Seventeen male WT CD-1 mice (63-86 days, *M* = 75.76 days, *SD* = 10.10), 13 female WT CD-1 mice (64-84 days, *M* = 76.23 days, *SD* = 7.97), 15 male BRL mice (72-77 days, *M* = 75.13 days, *SD* = 2.07), 14 female BRL mice (72-77 days, *M* = 75.57 days, *SD* = 2.21), five aged male Prrxl1^−/−^ mice (104-256 days, *M* = 219.60 days, *SD* = 65.06), six aged female Prrxl1^−/−^ mice (132-274 days, *M* = 216.67 days, *SD* = 69.58), 10 male Prrxl1^−/−^ mice (61-82 days, *M* = 68.90 days, *SD* = 7.31) and 11 female Prrxl1^−/−^ mice (64-96 days, *M* = 76.27 days, *SD* = 10.03) were used in the Textured Novel Object Recognition behavioral test (Table 1). A different set of twenty-seven male WT CD-1 mice (43-126 days, *M* = 107.00 days, *SD* = 15.79), 14 female WT CD-1 mice (46-121 days, *M* = 110.86 days, *SD* = 20.20), 15 male BRL mice (52-115 days, *M* = 109.20 days, *SD* = 15.89), 14 female BRL mice (52-117 days, *M* = 104.36 days, *SD* = 22.27), six male Prrxl1^−/−^ mice (43-106 days, *M* = 92.50 days, *SD* = 24.94) and eight female Prrxl1^−/−^ mice (46-127 days, *M* = 92.25 days, *SD* = 31.50) were used in the Open Field behavioral test (Table 2). Eight of the Prrxl1^−/−^ mice that were used in the Open Field test were reused for the aged group for the Textured Novel Object Recognition test. All of the mice were born, raised, and housed in the Queens College vivarium. The mice were maintained on a 12:12-hour light/dark schedule and given ad libitum access to food (5015 Mouse Diet, Lab Diet) and water.

**Table 1.** The strain, sex, count and average age of the mice used in the Textured Novel Object Recognition experiment.

| **Strain** | **Sex** | **Count** | **Average Age (days)** |
| --- | --- | --- | --- |
| WT | Male | 17 | 75.76 |
| WT | Female | 13 | 76.23 |
| BRL | Male | 15 | 75.13 |
| BRL | Female | 14 | 75.57 |
| Prrxl1<sup>-/-</sup> | Male | 5 | 219.60 |
| Prrxl1<sup>-/-</sup> | Female | 6 | 216.67 |
| Prrxl1<sup>-/-</sup> | Male | 10 | 68.90 |
| Prrxl1<sup>-/-</sup> | Female | 11 | 76.27 |

**Table 2.** The strain, sex, count and average age of the mice used in the Open Field experiment.

| **Strain** | **Sex** | **Count** | **Average Age (days)** |
| --- | --- | --- | --- |
| WT | Male | 27 | 107.00 |
| WT | Female | 14 | 110.86 |
| BRL | Male | 15 | 109.20 |
| BRL | Female | 14 | 104.36 |
| Prrxl1<sup>-/-</sup> | Male | 6 | 92.50 |
| Prrxl1<sup>-/-</sup> | Female | 8 | 92.25 |

### 2.2. Experimental Setup

#### 2.2.1. Behavior Arena Setup

The behavioral arena was constructed using four transparent acrylic sheets measuring 44 cm wide x 35 cm high, joined at the sides and attached to a base. A 0.08 cm gap was made between the bottom of one of the transparent acrylic sheets and the base to allow a 43 cm x 44 cm x 0.3 cm black acrylic sheet to slide in and out of the arena. A Monochrome Gig E camera (Basler) was centered 69.85 cm above the behavioral arena setup to record the activity. The camera was connected to a dedicated Dell desktop computer located in the same room, equipped with Pylon Viewer (Version 6.2.0.21487, 64-Bit) (Figure 1a). EthoVision XT Software (Version 17.0.1622) by Noldus was configured for the behavioral testing experiment. The behavioral arena was used for both the textured novel object recognition test and the open field test.

**Figure 1.**
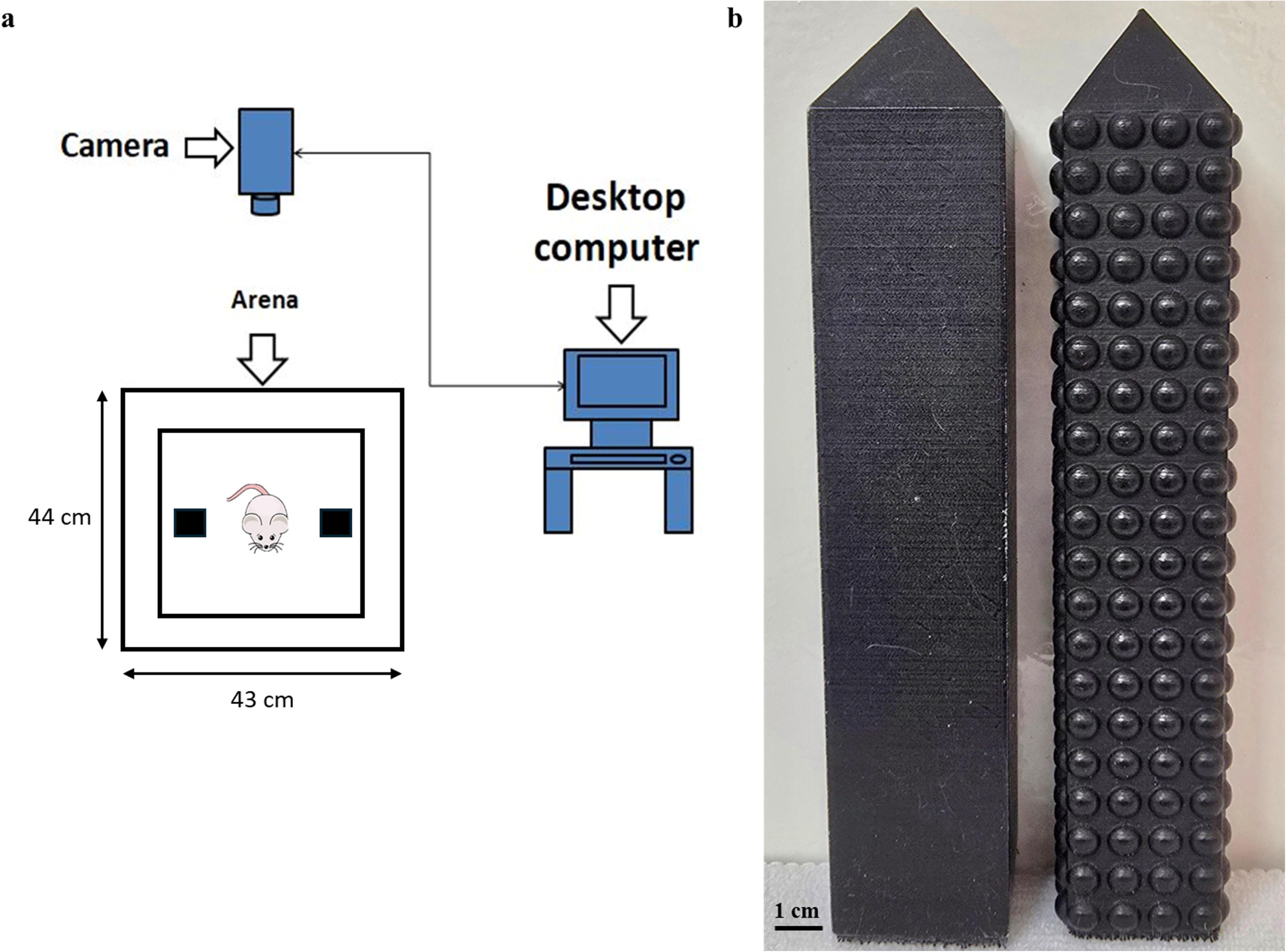
a) Cartoon showing the experiment setup, b) smooth/familiar (left) and uneven/novel (right) objects used in the experiment.

#### 2.2.2. Detection Settings

The detection setting in the EthoVision XT software was adjusted so that the mouse could be differentiated from the background. A non-experimental mouse was placed in the arena, and using the detection settings, a small rectangle was reshaped over the mouse in the arena to include the entire body and a small portion of the base of the tail. The pixels were adjusted from 64 to 225, which allowed for the mouse to be brighter compared to the background to aid in the subsequent tracking by the software.

### 2.3. Textured Novel Object Recognition Behavior Test

The novel object recognition test assesses learning and memory in rodents [14, 16]. In this study, the novel object recognition test has been adjusted to assess the ability to discriminate between two objects of different textures in a whisker-dependent manner. The textured novel object recognition test assesses the whisker-dependent discriminatory behavior of rodents [20, 21].

#### 2.3.1. Arena Setup

The objects used for this experiment were manufactured in the Queens College Makerspace. A smooth and uneven object was designed using the Tinkercad website. The smooth object was used as the familiar object, while the uneven object was used as the novel object (Figure 1b). The smooth object consisted of a pyramid measuring 3.30 cm x 3.30 cm x 1.96 cm placed on top of a rectangle measuring 3.30 cm x 3.30 cm x 14.90 cm for a total height of 16.85 cm. The uneven object consisted of a pyramid measuring 2.85 cm x 2.83 cm x 1.96 cm placed on top of a rectangle measuring 2.85 cm x 2.83 cm x 14.90 cm for a total height of 16.85 cm with eighty half-spheres measuring 0.23 cm x 0.60 cm placed 0.1 cm apart on each side of the rectangle. The measurement of the combined area of the base of the rectangle and the half-sphere of the uneven object was the same as the base of the smooth object. The design for the objects was exported as .STL files and then converted to a printable .ufp file using UltiMaker Cura Version 5.4.0. The objects were printed using an UltiMaker S3 printer using MatterHackers Build Series PLA Filament – 2.85mm (1kg), which was provided by the Queens College Makerspace. The objects were painted using non-toxic Rust-Oleum Painter’s Touch 2X Ultra Cover Spray Paint in Flat Black. The novel object testing occurred over two consecutive days. One object was placed 12.10 cm from the left edge, while the second object was placed 12.10 cm from the right edge of the acrylic sheet. The objects were placed 20.30 cm from the upper and lower edges of the acrylic sheet, with a distance of 12.20 cm between the two objects. The objects were attached to the acrylic sheet using Command Damage Free Hanging Strips, cut to size, and placed at the bottom of the objects and on the acrylic sheet. A 2 cm zone – the average length of the whiskers – was drawn around the objects. The object and the 2 cm zone around the object were combined into a cumulative zone that was used for analysis.

#### 2.3.2. Day 1 – Habituation

Two smooth objects were placed in the arena. The objects were adjusted so that they were located within the drawn outline. Each mouse was placed in the center of the arena and allowed to explore the arena for 10 minutes. EthoVision XT software began tracking the mouse when the nose point of the mouse was near any of the objects for ≥ 5.00s. At the end of each trial, each mouse was removed and returned to its cage. Each mouse was weighed before the start of the trial.

#### 2.3.3. Day 2 – Training Session and Testing Session

The habituation procedure conducted the previous day was replicated for the Training Session. The mouse spent 10 minutes in the arena with the two smooth objects. At the end of the trial, the mouse was removed and placed in its cage. After cleaning the objects with Dawn liquid soap and water, and the arena with a damp paper towel, the uneven object was added to the arena. The uneven object was counterbalanced, whether it was replaced by the smooth object on the right side or the left side of the arena. Roughly half of mice in each sex and genotype was exposed to the uneven object on the right side of the arena, while the other half was exposed to the object on the left side of the arena. The mouse was returned to the arena five minutes after being removed and spent 10 minutes exploring the familiar smooth object and the novel uneven object. At the end of each trial, the mouse was removed and returned to its cage. Each mouse was weighed before the start of the training session. The weight of the mice did not differ significantly between day 1 and day 2.

#### 2.3.4. Behavior Quantification

The time in seconds that the mouse spent in the arena began when the nose point of the mouse was in the 2 cm zone surrounding the objects for ≥ 5.00s. With the aid of video analysis, only the cumulative time in seconds that the mouse spent exploring the right familiar object and the left familiar object using the whiskers during the training session, and the cumulative time in seconds that the mouse spent exploring the familiar object and the novel object using the whiskers during the testing session was recorded. The velocity of each mouse was also recorded. The percentage of the time spent at the novel object was calculated using:

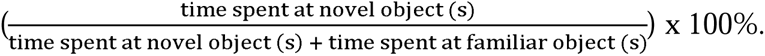

### 2.4. Open Field Behavior Test

The open field behavioral test was designed to assess anxiety levels, exploratory behavior, and emotionality in rats as they investigated an empty box-like structure [19]. Rodents with low anxiety levels will typically display increased exploratory behavior, greater locomotor activity, and an increase in time spent in the open inner area of the arena. On the other hand, rodents with high anxiety levels will tend to display decreased exploratory behavior, reduced locomotor activity, and an increase in time spent along the outer walls of the arena [17, 18].

#### 2.4.1. Arena Setup

The 43 cm x 44 cm x 0.3 cm black acrylic sheet was divided into an outer portion measuring 57.94 cm x 49.17 cm and an inner portion measuring 31.00 cm x 31.50 cm, each with an equal square area. The area of the outer arena alone measured 959.81 cm^2^, while the area of the inner arena measured 960.38 cm^2^.

#### 2.4.2. Day 1 Habituation & Day 2 Test

The open-field behavioral testing occurred over a period of two consecutive days, approximately 24 hours apart. The first day was the habituation period, while the second day was the testing period. On both days, a mouse was placed in the center of the arena and allowed to explore the box for 15 minutes. EthoVision XT software began tracking the mouse when the center point of the mouse was in the arena for ≥ 5.00s. The cumulative time in seconds that the mouse spent in the outer arena and the inner arena was recorded. The velocity of each mouse was also recorded. The percentage of time spent in the outer arena was calculated using:

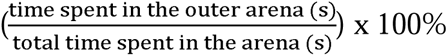. The velocity moved was calculated using: velocity 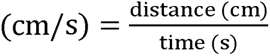.

### 2.5. Statistical Analyses

The data that was collected using the EthoVision XT software was organized and analyzed using the statistical program IBM SPSS (Version 29.0.0.0 (241)). One-sample *t*-tests were conducted to determine if there was a significant preference for either the right or left smooth object during the training phase. One-sample *t*-tests were conducted to determine if there was a significant preference for either the familiar smooth or the novel uneven object during the testing phase. Paired *t*-tests were conducted to compare the mean percentage of time spent in the outer arena to the inner arena for day 1 and day 2. Repeated-measures Analysis of Variances (ANOVA) were conducted to compare the mean time spent in the inner arena, and the mean velocity traveled on day 1 to day 2. Independent measures ANOVAs with Tukey’s HSD post-hoc were conducted to compare the mean time spent in the inner arena, the mean velocity traveled. Nested ANOVAs were conducted to examine sex differences within each of the three strains of mice. The statistical analyses were conducted using an α = .05 to determine significance.

Shapiro-Wilk Test was used to determine if there were any significant outliers. One from the WT group was removed from the Textured Novel Object Recognition analysis. Two from the BRL group and one from the WT group were removed from the Open Field analysis.

## 3. Results

### 3.1. Textured Novel Object Recognition Behavior Test

The novel object recognition behavior test is based on the observation that rodents prefer to explore new objects versus those they have encountered previously [16]. In the textured novel object recognition behavior paradigm, mice use their whiskers to discriminate between two different textures. The sensory information from the whiskers travels along the whisker-to-barrel cortex pathway to the somatosensory cortex for processing. During the training session, the mice showed no preference for the left or the right objects. During the testing session, we predicted that the mice would spend a greater percentage of the time investigating the novel texture. The mean percentage of time spent at the left object during the training session and the novel object during the testing session was compared to a test value of 50%. A preference for an object was demonstrated if the time spent at the object was greater than 50% of the total time spent investigating both objects. The minimum threshold for the exploration of both objects was 20 seconds [22].

#### 3.1.1. No preference for the novel object was displayed during the entire 10-minute duration

The mean percentage of time spent at the novel object was analyzed to determine if there was a significant preference for the novel object compared to the familiar object for the entire 10-minute duration (Figure 2a). The percentage of time spent at the novel object for male WT mice (*M* = 56.03%, *SD* = 15.75, *t*(16) = 1.58, *p* = .134), male BRL mice (*M* = 57.11%, *SD* = 15.95, *t*(14) = 1.73, *p* = .106), aged male Prrxl1^−/−^ mice (*M* = 59.58%, *SD* = 9.29, *t*(4) = 2.31, *p* = .082) and male Prrxl1^−/−^ mice (*M* = 54.58%, *SD* = 11.81, *t*(9) = 1.25, *p* = .242) were not significant. The percentage of time spent at the novel object for female WT mice (*M* = 52.23%, *SD* = 14.18, *t*(12) = 0.57, *p* = .581), female BRL mice (*M* = 52.66%, *SD* = 11.63, *t*(14) = 0.88, *p* = .391), aged female Prrxl1^−/−^ mice (*M* = 41.15%, *SD* = 11.46, *t*(5) = −1.89, *p* = .117) and female Prrxl1^−/−^ mice (*M* = 52.86%, *SD* = 20.73, *t*(10) = 0.46, *p* = .657) was also not significant. These findings suggest that none of the mice displayed a significant preference for either the novel or familiar objects during the entire 10-minute duration.

**Figure 2.**
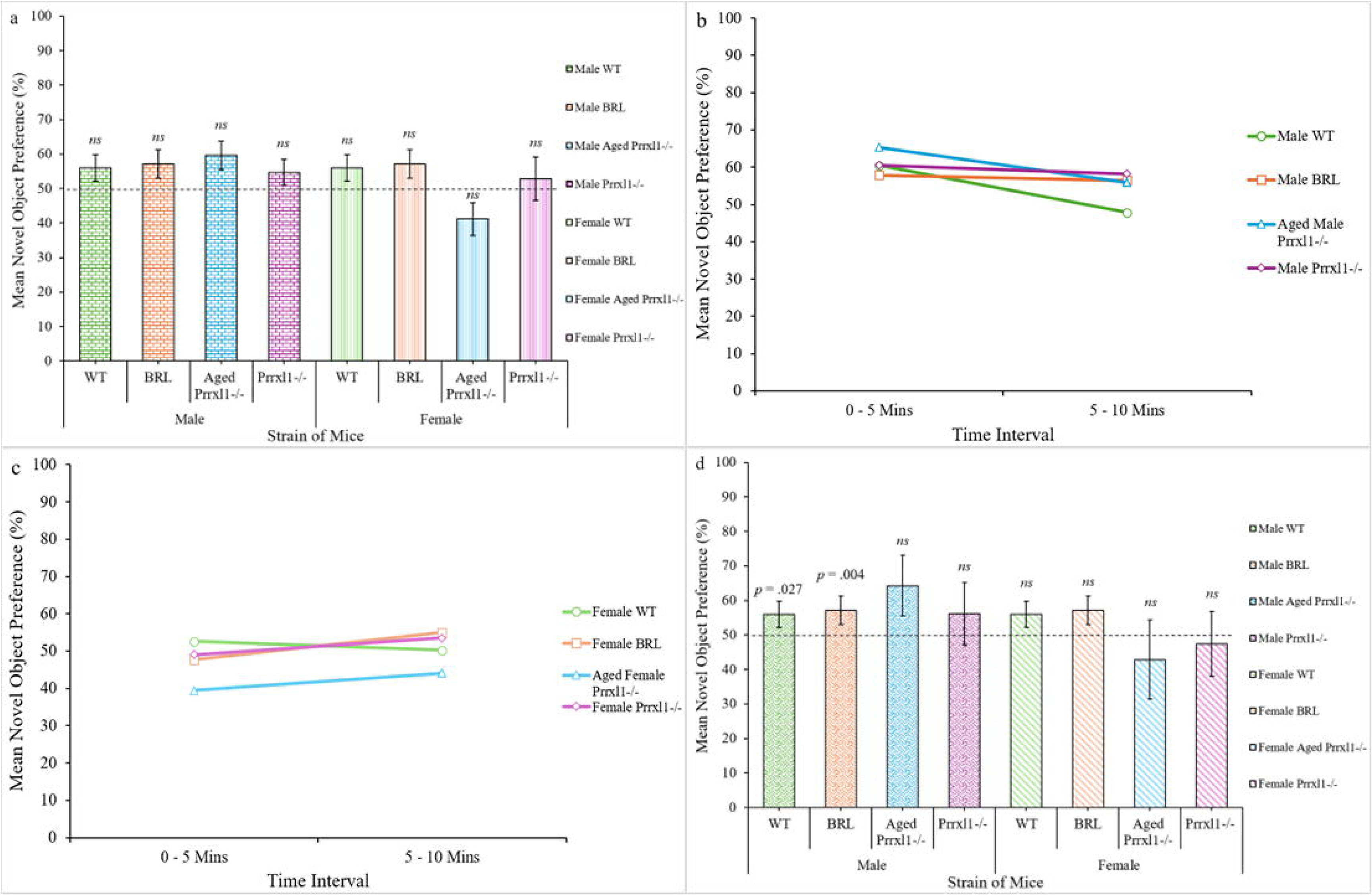
Graphs showing the mean novel object preference a) for all strains of mice during the entire 10-minute duration, b) across the two 5-minute intervals for the strains of male mice, c) across the two 5-minute intervals for the strains of female mice, and d) for all strains of mice during the first 3 minutes. The bars represent the population means; the error bars represent one standard error of the mean.

#### 3.1.2. Male mice and female WT mice displayed a decrease in the percentage of time spent at the novel object as time spent in the arena increased

We noticed the mice were less active during the second half of the 10-minute test period, so we evaluated performance during the first versus the second five-minute intervals. All strains of male mice spent more time exploring the novel object during the first 5 minutes compared to the second 5 minutes (Figure 2b). While female WT mice also spent more time exploring the novel object during the first 5 minutes (Figure 2c). Since the time spent investigating the novel object was greater during the first 5 minutes in the arena, the novel object preference during a shorter time – the first 3 minutes – was additionally analyzed. Additionally, there was no preference for the objects during the first 3 minutes of the training period.

#### 3.1.3. Male mice with no somatotopic distribution in the whisker-to-barrel cortex pathway and male mice with somatotopic disruption in the barrel cortex displayed a greater preference for the novel object

The mean percentage of time spent at the novel object was compared to a test value of 50% to determine if there was a significant preference for the novel object compared to the familiar object for the first 3 minutes (Figure 2d), an interval used in previous studies [23]. Male WT spent a significantly greater percentage of the time at the novel texture (*M* = 58.70%, *SD* = 14.70, *t*(16) = 2.44, *p* = .027). Male BRL mice also spent a significantly greater percentage of the time at the novel texture (*M* = 63.03%, *SD* = 14.94, *t*(14) = 3.38, *p* = .004). These findings suggest that male WT and male BRL mice were able to discriminate between the two textures using their whiskers. The percentage of time spent at the novel texture for aged male Prrxl1^−/−^mice (*M* = 64.39%, *SD* = 19.74, *t*(5) = 1.63, *p* = .179) and male Prrxl1^−/−^ mice (*M* = 56.15%, *SD* = 28.73, *t*(9) = 10.68, *p* = .515) were not significant. The percentage of time spent at the novel object for female WT mice (*M* = 57.62%, *SD* = 20.30, *t*(12) = 1.35, *p* = .201), female BRL mice (*M* = 53.30%, *SD* = 16.35, *t*(14) = 0.78, *p* = .447), aged female Prrxl1^−/−^ mice (*M* = 49.88%, *SD* = 28.08, *t*(5) = −0.62, *p* = .562) and female Prrxl1^−/−^ mice (*M* = 47.45%, *SD* = 30.99, *t*(10) = −0.27, *p* = .791) were also not significant. These findings suggest that male WT and male BRL mice were able to discriminate between the two textures using their whiskers, while both groups of male Prrxl1^−/−^ mice and all strains of female mice were unable to discriminate between the two textures using their whiskers during the first 3 minutes.

### 3.2. Open Field Behavior Test

The open field behavior test is used to study exploratory behavior and anxiety in rodents [24]. Mice, being prey animals, display an aversion to brightly lit, open areas. A willingness to explore the arena is associated with spending more time in the inner arena and greater velocity. On the other hand, less willingness to explore the arena is associated with staying closer to the walls in the outer arena, avoiding the “open area” of the inner arena, and slower velocity. In general, mice display less willingness to explore by spending more time in the outer arena compared to the inner arena, which represents the “open area”. We measured anxiety as a control behavior by analyzing the willingness or lack of willingness to explore the arena. In this study, we predicted that mice of all strains should also exhibit the same behavior, spending a greater percentage of the time in the outer arena compared to the inner arena.

#### 3.2.1. Male and female mice spent a significantly greater percentage of the time in the outer arena compared to the inner arena on days 1 and 2

Male WT mice spent 73.10% of the time in the outer arena on day 1, *t*(26) = 11.70, *p* <.001, and 82.50% of the time in the outer arena on day 2, *t*(26) = 18.31, *p* <.001. Male BRL mice spent 67.37% of the time in the outer arena on day 1, *t*(14) = 16.28, *p* <.001, and 76.00% of the time in the outer arena on day 2, *t*(14) = 16.20, *p* <.001. Male Prrxl1^−/−^ mice spent 73.56% of the time in the outer arena on day 1, *t*(5) = 3.24, *p* = .020, and 71.52% of the time in the outer arena on day 2, *t*(5) = 3.19, *p* = .020. Similarly, female WT mice spent 73.79% of the time in the outer arena on day 1, *t*(13) = 8.81, *p* <.001, and 79.54% of the time in the outer arena on day 2, *t*(13) = 13.16, *p* <.001. Female BRL mice spent 72.51% of the time in the outer arena on day 1, *t*(13) = 12.72, *p* <.001, and 73.52% of the time in the outer arena on day 2, *t*(13) = 11.81, *p* <.001. Female Prrxl1^−/−^ mice spent 71.00% of the time in the outer arena on day 1, *t*(7) = 5.27, *p* <.001, and 80.88% of the time in the outer arena on day 2, *t*(7) = 8.73, *p* <.001. These findings indicate that all mice, regardless of strain and sex, spent a significantly greater mean percentage of the time in the outer arena compared to the inner arena both on day 1 and day 2.

Heatmaps provide a visual representation of where the mouse spent the longest time in the arena, and were consistent with the time data. Merged heatmaps were plotted for the mean time the mice spent exploring the arena on day 1 (Figure 3a) and day 2 (Figure 3b). All mice spent more time in the outer arena compared to the inner arena, indicated by the presence of a lighter track around the arena edge and hot spots in the outer arena corners. In addition, the heatmaps contained more exploratory tracks in the inner arena on day 1 compared to day 2, indicating that the mice spent more time in the inner arena on day 1 compared to day 2. All mice, except for male Prrxl1^−/−^ mice, spent a greater percentage of time in the outer arena on day 2 compared to day 1.

**Figure 3.**
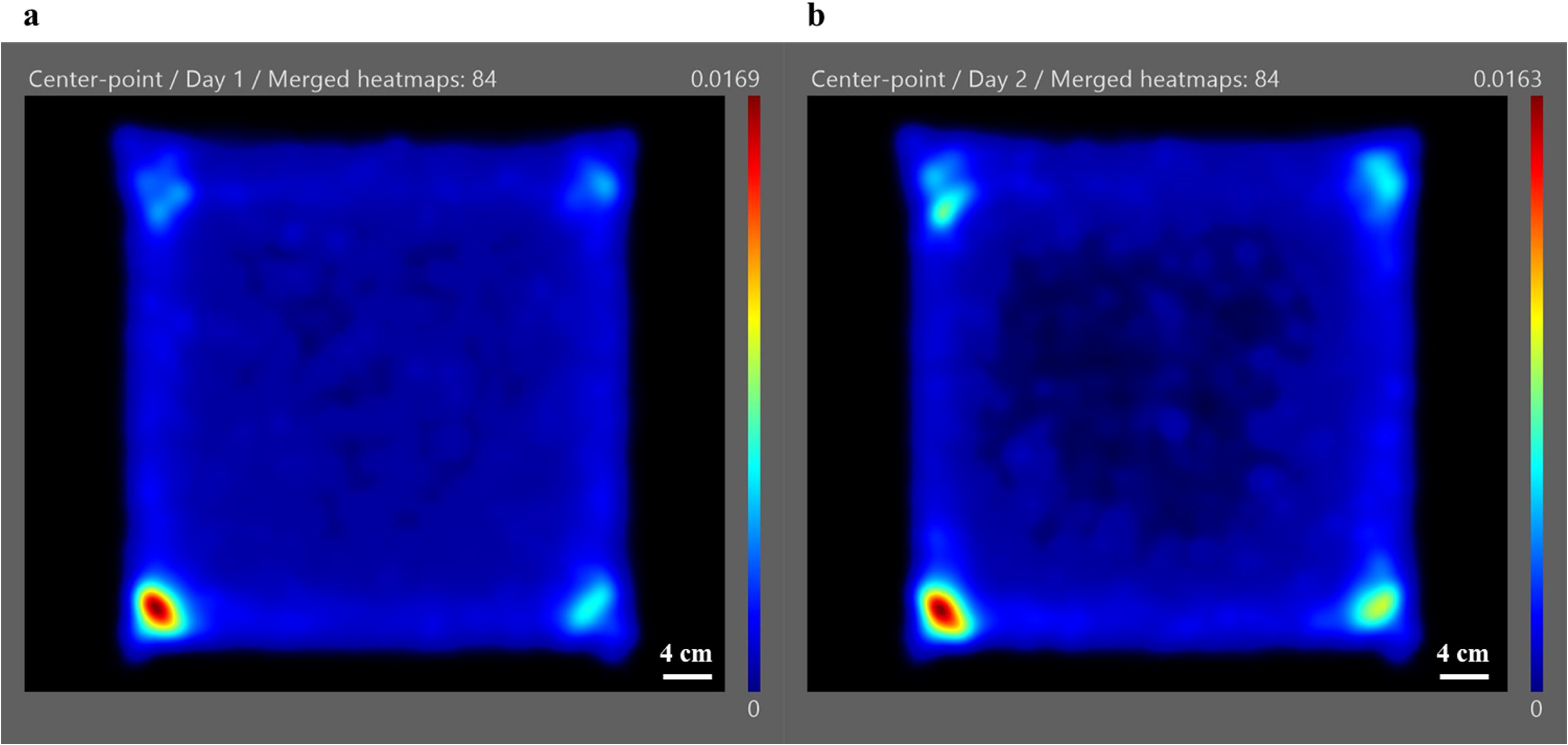
Merged heatmaps showing the mean time spent in the arena across a) day 1 and b) day 2. Hot spots - where the mouse spent a substantial amount of time - were red, while the color gradually changed to orange, then yellow, green, and finally to blue as the time spent in that area decreased.

#### 3.2.2. Male WT, male BRL, female WT, and female Prrxl1^−/−^ mice displayed significantly greater exploratory behavior on day 1 compared to day 2

The exploratory behavior – measured as time spent in the inner arena – of all strains of mice on day 1 was compared to day 2 (Figure 4a). Male WT mice spent significantly more time in the inner arena on day 1 (*M* = 241.90s, *SD* = 92.38) compared to day 2 (*M* = 157.46s, *SD* = 83.04), (*F*(1, 26) = 35.53, *p* <.001). Male BRL mice spent significantly more time in the inner arena on day 1 (*M* = 293.69s, *SD* = 37.19) compared to day 2 (*M* = 216.02s, *SD* = 55.95), (*F*(1, 14) = 43.31, *p* <.001). There was no significant difference between the time spent in the inner arena on day 1 (*M* = 238.03s, *SD* = 160.34) compared to day 2 (*M* = 256.37s, *SD* = 148.93) for male Prrxl1^−/−^ mice (*F*(1, 5) = 0.11, *p* = .758).

**Figure 4.**
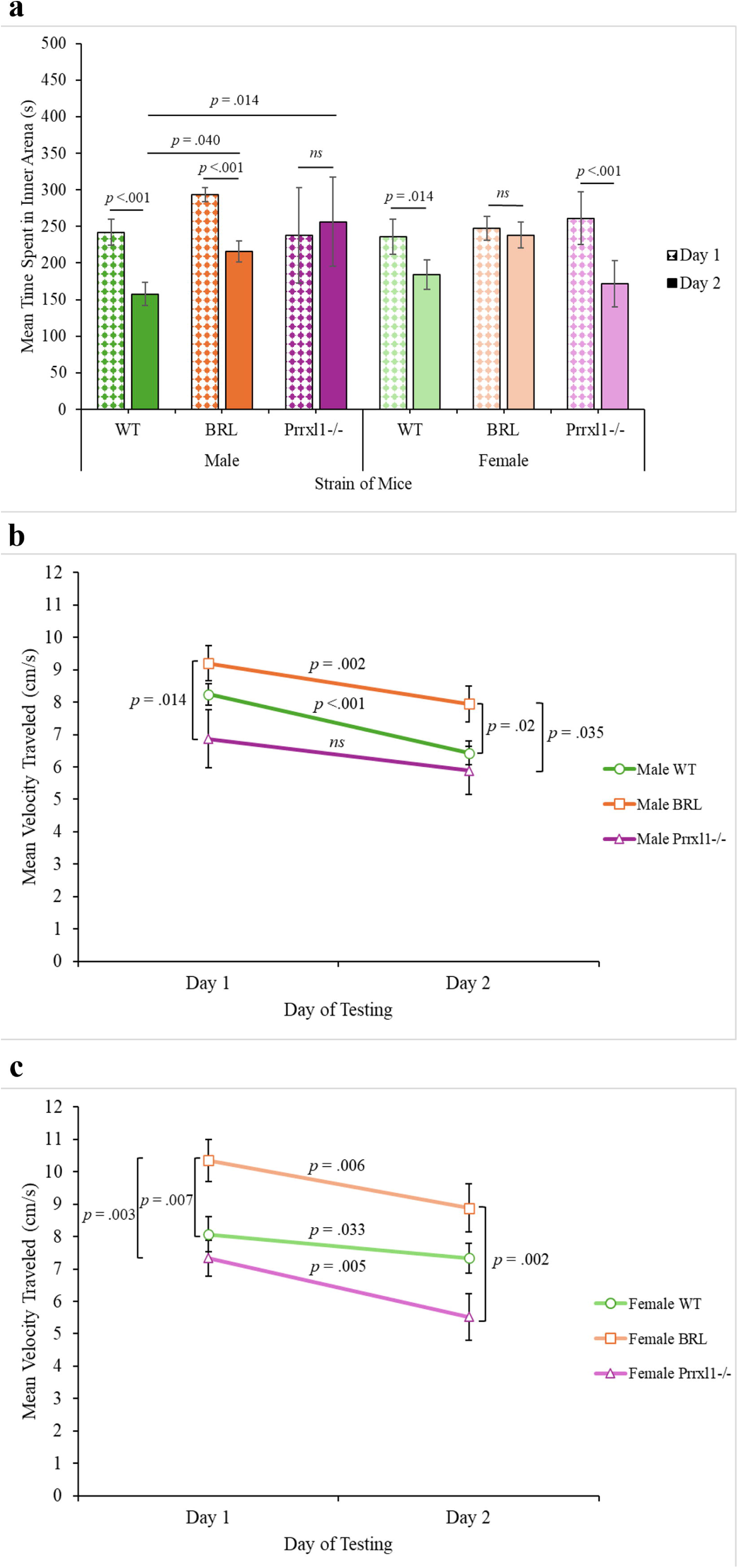
Graphs showing a) the mean time spent in the inner arena for the three strains of male and female mice, b) the mean velocity traveled for the three strains of male mice, and c) the mean velocity traveled for the three strains of female mice. The bars represent the population means; the error bars represent one standard error of the mean.

Female WT mice spent significantly more time in the inner arena on day 1 (*M* = 235.91s, *SD* = 90.90) compared to day 2 (*M* = 184.14s, *SD* = 75.61), (*F*(1, 13) = 9.17, *p* = .010). Female Prrxl1^−/−^ mice spent significantly more time in the inner arena on day 1 (*M* = 260.98s, *SD* = 101.47) compared to day 2 (*M* = 171.88s, *SD* = 89.97), (*F*(1, 7) = 31.75, *p* <.001). There was no significant difference between the time spent in the inner arena on day 1 (*M* = 247.36s, *SD* = 59.57) compared to day 2 (*M* = 238.18s, *SD* = 67.14) for female BRL mice (*F*(1, 13) = 0.32, *p* = .581). Taken together, these findings suggest that male and female WT mice, male BRL mice, and female Prrxl1^−/−^ mice displayed greater exploratory behavior on day 1 compared to day 2 by spending more time in the inner arena.

#### 3.2.3. Male mice with somatotopic disruption in the whisker-to-barrel cortex pathway displayed significantly greater exploratory behavior than male WT mice on day 2

Exploratory behavior was compared between strains separately for males and females. (Figure 4a). Within males on day 1, there was no significant difference between the mean time spent in the inner arena among WT mice (*M* = 241.90s, *SD* = 92.38), BRL mice (*M* = 293.69s, *SD* = 37.19), and Prrxl1^−/−^ mice (*M* = 238.03s, *SD* = 160.34), (*F*(2, 45) = 1.73, *p* = .188). In contrast, within males on day 2, BRL mice (*M* = 216.02s, *SD* = 55.95) and Prrxl1^−/−^ mice (*M* = 256.37s, *SD* = 148.93) displayed greater exploratory behavior by spending significantly more time in the inner arena compared to WT mice (*M* = 157.46s, *SD* = 83.04), (*F*(2, 45) = 4.38, *p* = .018). These findings suggest that male BRL and male Prrxl1^−/−^ mice displayed greater exploratory behavior compared to male WT mice on day 2.

However, within females on day 1, there was no significant difference between the mean time spent in the inner arena among WT mice (*M* = 235.91s, *SD* = 90.90), BRL mice (*M* = 247.36s, *SD* = 59.57), and Prrxl1^−/−^ mice (*M* = 260.98s, *SD* = 101.47), (*F*(2, 33) = 0.24, *p* = .790). Similarly, within females on day 2, there was no significant difference between the mean time spent in the inner arena among WT mice (*M* = 184.14s, *SD* = 75.61), BRL mice (*M* = 238.18s, *SD* = 67.14), and Prrxl1^−/−^ mice (*M* = 171.88s, *SD* = 89.97), (*F*(2, 33) = 2.62, *p* = .088).

#### 3.2.4. Male BRL mice moved significantly faster than male Prrxl1^−/−^ mice on day 1 and day 2, and male WT mice on day 2

Male WT mice moved significantly faster on day 1 (*M* = 8.25 cm/s, *SD* = 1.71) compared to day 2 (*M* = 6.43 cm/s, *SD* = 1.87), (*F*(1, 26) = 30.05, *p* <.001). Male BRL mice also moved significantly faster on day 1 (*M* = 9.20 cm/s, *SD* = 2.09) compared to day 2 (*M* = 7.95 cm/s, *SD* = 2.15), (*F*(1, 14) = 13.65, *p* = .002). Although male Prrxl1^−/−^ mice moved faster on day 1 (*M* = 6.87 cm/s, *SD* = 2.21) compared to day 2 (*M* = 5.89 cm/s, *SD* = 1.79), this difference was not significant, (*F*(1, 5) = 3.42, *p* = .124). On day 1, male BRL mice moved significantly faster than male Prrxl1^−/−^ mice, (*F*(2, 45) = 3.39, *p* = .042). On day 2, male BRL mice moved significantly faster than male WT mice, and male Prrxl1^−/−^ mice, (*F*(2, 45) = 3.70, *p* = .033). These findings suggest that male BRL mice displayed greater exploratory behavior by moving faster during the time spent in the arena.

#### 3.2.5. Female BRL moved significantly faster than female Prrxl1^−/−^ mice on day 1 and day 2, and female WT mice on day 2

Female WT mice moved significantly faster on day 1 (*M* = 8.07 cm/s, *SD* = 2.02) compared to day 2 (*M* = 7.34 cm/s, *SD* = 1.74), (*F*(1, 13) = 5.66, *p* = .033). Female BRL mice also moved significantly faster on day 1 (*M* = 10.35 cm/s, *SD* = 2.42) compared to day 2 (*M* = 8.88 cm/s, *SD* = 2.75), (*F*(1, 13) = 10.50, *p* = .006). Female Prrxl1^−/−^ mice also moved significantly faster on day 1 (*M* = 7.33 cm/s, *SD* = 1.59) compared to day 2 (*M* = 5.52 cm/s, *SD* = 2.03), (*F*(1, 7) = 16.60, *p* = .005). On day 1, female BRL mice moved significantly faster than female WT mice and female Prrxl1^−/−^ mice, (*F*(2, 33) = 6.58, *p* = .004). On day 2, female BRL mice moved significantly faster than female Prrxl1^−/−^ mice, (*F*(2, 33) = 5.78, *p* = .007). These findings suggest that female BRL mice displayed greater exploratory behavior by moving faster during the time spent in the arena.

## 4. Discussion

We investigated the impact of somatotopic pattern disruption on textured novel object recognition and open field behavioral testing and found that the disruption in the whisker-to-barrel pathway alters behavior. In the textured novel object recognition behavioral test, only the male BRL mice were able to discriminate between the textures, and did so during the first 3 minutes of object exposure. Male BRL also spent more time exploring in the open field behavioral test than did male WT and Prrxl1^−/−^ mice. Both male and female BRL mice moved faster across both days of the open field behavioral test.

### 4.1. Textured Novel Object Recognition Behavioral Test

The textured novel object recognition task tests the ability to differentiate between a familiar texture and a novel texture using their whiskers [14, 20, 21, 25]. Mice rely on their whiskers rather than their visual system to explore their environment [1, 2, 26]. Using the somatosensory system is necessary when testing the CD-1 strain, which has poor eyesight [27]. Therefore, visual cues were not sufficient to discriminate between the novel and familiar objects, nor were olfactory cues, since the two objects were made of the same material and thoroughly cleaned between trials. The main difference between these two objects was that the familiar object was smooth, while the novel object consisted of bumps made of the same material, which gave it an uneven texture; thus, the mice used their whisker to differentiate between the novel and familiar textures. The mice were able to successfully discriminate between the novel texture and the familiar texture if the percentage of time spent investigating the novel object was significantly greater compared to the familiar object. This result also suggests that the short-term memory of the mice was intact. A time span of 5 minutes was used between the training session and the testing session to minimize hippocampal-mediated learning [21, 25, 28].

Although no mice spent a greater percentage of the time exploring the novel object rather than the familiar object for the entire 10-minute test period, we observed that all the strains of male mice and female WT mice spent less time investigating the novel textures as the time spent in the arena increased. One possible explanation is that the mice eventually lost interest in the novel texture, since it was no longer “new” resulting in no preference. This was supported by comparing the first half to the second half of the 10-minute test period, during which the preference for the novel texture decreased as the time spent in the arena increased. Based on the interval used in Chen et al., [23], we analyzed the first 3 minutes and found that male WT mice and male BRL mice spent a significantly greater percentage of the time investigating the novel texture, which demonstrated the ability to discriminate between the two textures.

When sex differences were compared, we found that all the strains of male mice and female WT and BRL mice demonstrated a preference for the novel texture by spending more time investigating the novel texture. However, the aged female Prrxl1^−/−^ mice demonstrated a preference for the familiar texture by spending more time investigating the familiar texture. Additionally, only male WT and male BRL mice displayed a significant preference for the novel texture during the first 3 minutes. In a previous study, Arakawa et al., [7] reported that BRL mice were unable to discriminate between textures in a whisker-dependent manner. However, unlike our methodology, a time span of 1 hour was used between the training session and the testing session, possibly resulting in the inability to remember the previously encountered familiar texture during the testing session. Additionally, the objects used were prone to odor absorption and were very visually different, confounds that were largely avoided in our paradigm.

Overall, we found that male mice with somatotopic disruption in the barrel cortex displayed a significantly greater preference for the novel texture, indicating that their ability to discriminate between the two textures in a whisker-dependent manner was intact. However, the ability to discriminate was not found in female mice. The finding that males spent more time exploring the novel object was not corroborated by previous research, which demonstrated that adult female rodents spent significantly more time than adult male rodents exploring the novel object [29, 30].

### 4.2. Open Field Behavioral Test

The open field behavioral test was used to investigate the exploratory behavior of the mice by measuring the percentage of time spent in the outer arena compared to the inner arena. We observed that mice prefer to spend more time in the outer arena along the walls compared to the open inner arena, even after being habituated. These results support previous studies in which mice spent more time in the outer arena and less time in the inner arena [17–19, 24, 31]. We observed that all strains and sexes of mice, except for male Prrxl1^−/−^ mice, spent a greater percentage of the time in the outer arena on day 2 compared to day 1, indicating a decrease in exploratory behavior after becoming habituated to the arena. Additionally, male BRL mice and male Prrxl1^−/−^ mice spent significantly less time in the outer arena on day 2 compared to male WT mice, indicating greater exploratory behavior. We also investigated the velocity traveled across both days and observed that all mice moved faster on day 1 compared to day 2, suggesting that the mice displayed less exploratory behavior as the time spent in the arena increased. When sex differences were analyzed, male BRL mice and female BRL mice exhibited greater exploratory behavior among the strains by traveling a greater distance, while male Prrxl1^−/−^ mice and female Prrxl1^−/−^ mice displayed lower exploratory behavior.

Due to the initial limited availability of Prrxl1^−/−^ mice, some of the Prrxl1^−/−^ mice used in the open field experiment were also used for the textured novel object recognition experiment, meaning that some of the Prrxl1^−/−^ mice were not naive. Unfortunately, Prrxl1^−/−^ mice frequently have short lifespans, resulting in a decrease in the availability of Prrxl1^−/−^ mice [11]. The Prrxl1^−/−^mice that were reused were grouped with older Prrxl1^−/−^ mice and although the possibility exists that the increased age of the Prrxl1^−/−^ mice may play a factor in the ability to discriminate, the decrease in the ability to discriminate was also seen in the younger Prrxl1^−/−^ mice.

## 5. Conclusion

We observed that BRL mice in which the somatotopic representation is abolished in the barrel cortex exhibited greater exploratory behavior and were able to discriminate between textures in a whisker-dependent manner. On the other hand, Prrxl1^−/−^ mice in which the somatotopic representation is abolished in the PrV of the lemniscal brainstem nucleus exhibited less exploratory behavior and were unable to discriminate between textures. Our observations suggest that alterations in the somatotopic patterning in the whisker-to-barrel cortex pathway have an effect on the whisker-dependent discriminatory behavior in mice.

## Acknowledgments

This work was funded by grants NS126987 and GM122657 to JCB.

The authors would like to thank Dr. Carolyn L. Pytte for her insightful comments on our manuscript.

## Author contributions: CRediT

**Joanna M.F. Lutchman**: Conceptualization, Methodology, Formal Analysis, Investigation, Writing – Original Draft, Visualization.

**Sarah A.F. Lutchman**: Conceptualization, Methodology, Formal Analysis, Writing – Review & Editing.

**Joshua C. Brumberg**: Conceptualization, Methodology, Resources, Writing – Review & Editing, Supervision, Funding Acquisition.

## References

[1] Adibi, M. Whisker-Mediated Touch System in Rodents: From Neuron to Behavior. Frontiers in Systems Neuroscience 13 (2019): 40. 10.3389/fnsys.2019.00040

[2] Petersen, C. C. H. Sensorimotor processing in the rodent barrel cortex. Nature Reviews Neuroscience 20 (2019): 533–546. 10.1038/s41583-019-0200-y

[3] Li, H., & Crair, M. C. How do barrels form in somatosensory cortex? Annals of the New York Academy of Sciences 1225, no. 1 (2011): 119–129. 10.1111/j.1749-6632.2011.06024.x

[4] Woolsey, T. A., & Van Der Loos, H. The structural organization of layer IV in the somatosensory region (S I) of mouse cerebral cortex. Brain Research 17, no. 2 (1970): 205–242. 10.1016/0006-8993(70)90079-X

[5] Abdel-Majid, R. M., Leong, W. L., Schalkwyk, L. C., Smallman, D. S., Wong, S. T., Storm, D. R., Fine, A., Dobson, M. J., Guernsey, D. L., & Neumann, P. E. Loss of adenylyl cyclase I activity disrupts patterning of mouse somatosensory cortex. Nature Genetics 19 (1998): 289–291. 10.1038/980

[6] Welker, E., Armstrong-James, M., Bronchti, G., Ourednik, W., Gheorghita-Baechler, F., Dubois, R., Guernsey, D. L., Van Der Loos, H., & Neumann, P. E. Altered Sensory Processing in the Somatosensory Cortex of the Mouse Mutant Barrelless. Science 271, no. 5257 (1996): 1864–1867. 10.1126/science.271.5257.1864

[7] Arakawa, H., Akkentli, F., & Erzurumlu, R. S. Region-Specific Disruption of Adenylate Cyclase Type 1 Gene Differentially Affects Somatosensorimotor Behaviors in Mice. Eneuro 1, no. 1 (2014) ENEURO.0007-14.2014. 10.1523/ENEURO.0007-14.2014

[8] Iwasato, T., Inan, M., Kanki, H., Erzurumlu, R. S., Itohara, S., & Crair, M. C. Cortical Adenylyl Cyclase 1 Is Required for Thalamocortical Synapse Maturation and Aspects of Layer IV Barrel Development. Journal of Neuroscience 28, no. 23 (2008): 5931–5943. 10.1523/JNEUROSCI.0815-08.2008

[9] Chen, Z.-F., Rebelo, S., White, F., Malmberg, A. B., Baba, H., Lima, D., Woolf, C. J., Basbaum, A. I., & Anderson, D. J. The Paired Homeodomain Protein DRG11 Is Required for the Projection of Cutaneous Sensory Afferent Fibers to the Dorsal Spinal Cord. Neuron 31, no. 1 (2001): 59–73. 10.1016/S0896-6273(01)00341-5

[10] Ding, Y.-Q., Yin, J., Xu, H.-M., Jacquin, M. F., & Chen, Z.-F. Formation of Whisker-Related Principal Sensory Nucleus-Based Lemniscal Pathway Requires a Paired Homeodomain Transcription Factor, *Drg11*. The Journal of Neuroscience 23, no. 19 (2003): 7246–7254. 10.1523/JNEUROSCI.23-19-07246.2003

[11] Bakalar, D., Tamaiev, J., Zeigler, H. P., & Feinstein, P. Abolition of lemniscal barrellette patterning in *Prrxl1* knockout mice: Effects upon ingestive behavior. Somatosensory & Motor Research 32, no. 4 (2015): 236–248. 10.3109/08990220.2015.1086327

[12] Jacquin, M. F., Arends, J. J. A., Xiang, C., Shapiro, L. A., Ribak, C. E., & Chen, Z.-F. In *DRG11* Knock-Out Mice, Trigeminal Cell Death Is Extensive and Does Not Account for Failed Brainstem Patterning. The Journal of Neuroscience 28, no. 14 (2008): 3577–3585. 10.1523/JNEUROSCI.4203-07.2008

[13] Monteiro, C., Dourado, M., Matos, M., Duarte, I., Lamas, S., Galhardo, V., & Lima, D. Critical care and survival of fragile animals: The case of Prrxl1 knockout mice. Applied Animal Behaviour Science 158 (2014): 86–94. 10.1016/j.applanim.2014.06.007

[14] Antunes, M., & Biala, G. The novel object recognition memory: Neurobiology, test procedure, and its modifications. Cognitive Processing 13 (2012): 93–110. 10.1007/s10339-011-0430-z

[15] Ennaceur, A. One-trial object recognition in rats and mice: Methodological and theoretical issues. Behavioural Brain Research 215, no. 2 (2010): 244–254. 10.1016/j.bbr.2009.12.036

[16] Ennaceur, A., & Delacour, J. A new one-trial test for neurobiological studies of memory in rats. 1: Behavioral data. Behavioural Brain Research 31, no. 1 (1988): 47–59. 10.1016/0166-4328(88)90157-X

[17] Ennaceur, A. Tests of unconditioned anxiety—Pitfalls and disappointments. Physiology & Behavior 135 (2014): 55–71. 10.1016/j.physbeh.2014.05.032

[18] Gould, T. D., Dao, D. T., & Kovacsics, C. E. The Open Field Test. In T. D. Gould (Ed.), Mood and Anxiety Related Phenotypes in Mice. Neuromethods 42 (2009): 1–20. Humana Press. 10.1007/978-1-60761-303-9_1

[19] Hall, C. S. Emotional behavior in the rat. I. Defecation and urination as measures of individual differences in emotionality. Journal of Comparative Psychology 18, no. 3 (1934): 385–403. 10.1037/h0071444

[20] Hayashi, Y., Alamir, N., Sun, G., Tamagnini, F., Hayashi, Y., Williams, C., & Zheng, Y. An effective textured Novel Object Recognition Test (tNORT) for repeated measure of whisker sensitivity of rodents. Behavioural Brain Research 472 (2024): 115153. 10.1016/j.bbr.2024.115153

[21] Mattioni, L., Barbieri, A., Grigoli, A., Balasco, L., Bozzi, Y., & Provenzano, G. Alterations of Perineuronal Net Expression and Abnormal Social Behavior and Whisker-dependent Texture Discrimination in Mice Lacking the Autism Candidate Gene Engrailed 2. Neuroscience 546 (2024): 63–74. 10.1016/j.neuroscience.2024.03.023

[22] Lueptow, L. M. Novel Object Recognition Test for the Investigation of Learning and Memory in Mice. Journal of Visualized Experiments 126 (2017): e55718. 10.3791/55718

[23] Chen, C.-C., Lu, J., Yang, R., Ding, J. B., & Zuo, Y. Selective activation of parvalbumin interneurons prevents stress-induced synapse loss and perceptual defects. Molecular Psychiatry 23 (2018): 1614–1625. 10.1038/mp.2017.159

[24] Seibenhener, M. L., & Wooten, M. C. Use of the Open Field Maze to Measure Locomotor and Anxiety-like Behavior in Mice. Journal of Visualized Experiments 96 (2015): e52434. 10.3791/52434

[25] Wu, H.-P. P., Ioffe, J. C., Iverson, M. M., Boon, J. M., & Dyck, R. H. Novel, whisker-dependent texture discrimination task for mice. Behavioural Brain Research 237 (2013): 238–242. 10.1016/j.bbr.2012.09.044

[26] Gillespie, D., Yap, M. H., Hewitt, B. M., Driscoll, H., Simanaviciute, U., Hodson-Tole, E. F., & Grant, R. A. Description and validation of the LocoWhisk system: Quantifying rodent exploratory, sensory and motor behaviours. Journal of Neuroscience Methods 328 (2019): 108440. 10.1016/j.jneumeth.2019.108440

[27] Abdeljalil, J., Hamid, M., Abdel-mouttalib, O., Stéphane, R., Raymond, R., Johan, A., José, S., Pierre, C., & Serge, P. The optomotor response: A robust first-line visual screening method for mice. Vision Research 45, no. 11 (2005): 1439–1446. 10.1016/j.visres.2004.12.015

[28] Balasco, L., Pagani, M., Pangrazzi, L., Chelini, G., Ciancone Chama, A. G., Shlosman, E., Mattioni, L., Galbusera, A., Iurilli, G., Provenzano, G., Gozzi, A., & Bozzi, Y. Abnormal whisker-dependent behaviors and altered cortico-hippocampal connectivity in *Shank3b* ^−/−^ mice. Cerebral Cortex 32, no. 14 (2022): 3042–3056. 10.1093/cercor/bhab399

[29] Bettis, T., & Jacobs, L. F. Sex differences in object recognition are modulated by object similarity. Behavioural Brain Research 233, no. 2 (2012): 288–292. 10.1016/j.bbr.2012.04.028

[30] Cyrenne, D. M., & Brown, G. R. Ontogeny of sex differences in response to novel objects from adolescence to adulthood in lister hooded rats. Developmental Psychobiology 53, no. 7 (2011): 670–676. 10.1002/dev.20542

[31] Prut, L., & Belzung, C. The open field as a paradigm to measure the effects of drugs on anxiety-like behaviors: A review. European Journal of Pharmacology 463, no. 1-3 (2003): 3–33. 10.1016/S0014-2999(03)01272-X

